# Validating the Target Functions and Synergistic Multi-Target, Multi-Pathway Action Mode of Compound Kushen Injection Using CRISPR/CAS

**DOI:** 10.1101/2024.05.03.592304

**Authors:** Hanyuan Shen, Saeed Nourmohammadi, Yan Zhou, Yuka Harata-Lee, Zhipeng Qu, Wei Wang, Andrea J Yool, David L. Adelson

## Abstract

**Background and Purpose:** Due to the complexity of traditional Chinese medicine based on complex mixtures of natural products and their multi-target mechanism of action, the discovery and validation of relevant targets have always been challenging. In previous studies, using transcriptomic methods and Compound Kushen Injection (CKI) as a model drug, we identified multiple pathways and target genes through which CKI exerts its pharmacological effects. Therefore, we wished to verify these targets by perturbing those genetic pathways.

**Experimental Approach:** In this study, we selected eight key genes from four candidate pathways and used CRISPR/CAS technology to knock out these genes in four cell lines, validating their role in CKI activity.

**Key Results:** Firstly, we found that although the sensitivity of different cell lines to gene knockout varied, overall, it led to a reduction in various cellular activities. After the addition of CKI, we observed that, except for the minor impact of CDKN1A gene knockout on the effect of CKI, knocking out the other genes significantly affected the pharmacological efficacy of CKI in different assays. Among them, knockout of MYD88 and NFkB genes enhanced the efficacy of CKI. At the same time, we found that the genes IL24 and CYP1B1 play a crucial role in CKI inhibition of tumour cell migration, and the CYP1A1 gene is critical for the cell cycle arrest induced by CKI.

**Conclusion and Implications:** These find**ing**s not only validate the results of our previous transcriptomic analysis but also further demonstrate the complexity of pharmacological mechanisms of multi-target synergistic action of natural product mixtures.

**What is already known:** CKI demonstrates antitumor effects in both clinical and pharmacological research. Transcriptomic analysis shows CKI can perturb the expression of numerous genes in pathways related to cancer.

**What does this study add:** The knockout of most selected genes whose expression is altered by CKI can significantly affect the pharmacological effects of CKI.

IL24 and CYP1B1genes are essential to CKI’s inhibition of cancer cell migration and CYP1A1 is important for CKI’s G2M cell cycle arrest effect.

**What is the clinical significance:** The efficacy of CKI is demonstrated to arise from the synergistic action of multiple pathways and targets.

## 1 Introduction

Compound Kushen Injection (CKI), also known as Yanshu injection, is derived from the herbs Kushen (Radix *Sophorae flavescentis*) and Baituling (Rhizoma *Smilacis glabrae*)^1^ and is a modern Chinese herbal preparation based on traditional principles. CKI was approved by the China Food and Drug Administration in 1995 and has been used in clinical treatments for various types of cancer^2^. Despite substantial clinical evidence supporting the efficacy of CKI, the specific mechanisms of its action have remained unclear due to the complexity of its traditional Chinese medicine formulation. In our previous studies, we used CKI as a model traditional Chinese medicine formulation. We employed systems biology and functional genomics methods to investigate its potential anticancer mechanisms^3,4^. Our findings indicated that CKI can influence numerous genes and pathways associated with cancer treatment^5,6^. These results suggest that CKI operates through a multi-target, synergistic approach, which is a distinctive feature of traditional Chinese medicines. However, while we have confirmed the perturbation of specific genes and their corresponding protein products by CKI, most of the genes and pathways we identified still required further validation.

Based on our previous transcriptome results, we identified four pathways that are significantly affected by CKI and closely related to the tumorigenesis process. Subsequently, we selected two target genes within each pathway for knockout to validate their role. These selected genes exhibited substantial changes in their expression levels upon treatment with CKI, and they are considered core or essential nodes within their respective pathways. The genes we focused on are as follows:

Cell Cycle Pathway: CDKN1A, CDC14B

Cytokine-Cytokine Receptor Interaction: IL1A, IL24

NF-κB Signaling Pathway: MYD88, NFKB

Metabolism of Xenobiotics by Cytochrome P450 Pathway: CYP1A1, CYP1B1

It’s worth noting that, apart from MYD88, all the other genes exhibited significant upregulation upon treatment with CKI in the MDA-MB-231 and HepG2 cell lines (Supplementary Figure 1). This indicates that CKI has a substantial impact on the expression of these genes, particularly in the context of these cancer cell lines.

In the field of cancer biology, the CRISPR/CAS technique is widely employed to explore the genetic underpinnings of tumour initiation, progression, and therapeutic responses^7,8^. In addition, CRISPR/CAS has been used to investigate drug resistance in various cancer types, including those subjected to chemotherapy, targeted therapies, and inhibitors^9,10^. In this research project, we applied the CRISPR/CAS technique to validate the significance of genes and pathways involved in CKI’s anticancer properties. We conducted individual knockouts of the 8 selected genes in four distinct cell lines and observed the resulting phenotypic changes and their responses to CKI treatment. The findings revealed notable differences among the various cell lines. However, as a general trend, the knockout of these genes predominantly inhibited cellular functions and the effects of CKI. These results provide further validation for the data obtained from our previous transcriptomic analysis. They underscore that CKI’s anticancer effects are achieved through a synergistic action that perturbs multiple pathways and targets, solidifying our understanding of CKI’s mechanism of action in cancer treatment.

## 2 Methods

### 2.1 Cell culturing conditions

The MDA-MB-231 and B16F1 cells were cultured in DMEM medium supplemented with 10% fetal bovine serum (FBS). The A431 and E0771 cells were cultured in RPMI-1640 medium supplemented with 10% FBS. All cells were incubated at 37°C in a humidified incubator with 5% carbon dioxide (CO2).

### 2.2 Plasmid preparation

In the database of https://www.benchling.com/, based on the information of each target gene, guide sequences were designed on both sides of the selected exons. Short DNA oligonucleotides encoding the guide sequences were synthesized by Sigma. The synthesized oligonucleotides were annealed in the reaction system and ligated into the target plasmid through a one-step digestion-ligation reaction^11^. The newly synthesized plasmids were transfected into DH5-alpha competent cells (Thermo Fisher Scientific, MA, USA), and the transfected cells were plated on LB agar plates containing ampicillin for selection. One or two colonies (white, round-shaped) were picked and inoculated in LB liquid medium for overnight culture and amplification. The next day, plasmids were extracted from the competent cells with QIAprep Spin Miniprep Kit (Qiagen, Venlo, Netherlands) and sent to the Australian Genome Research Facility for Sanger sequencing to verify the correct insertion of the guide sequences. The verified plasmids were stored at -20 degrees Celsius.

### 2.3 Transfection and culturing of knockout cells

For plasmid transfection, we used the Neon Transfection System from Thermo Fisher Scientific. Prior to the actual transfection, preliminary experiments were conducted for each cell line to determine the optimal electroporation parameters, including voltage, pulse width, and pulse number. The transfection efficiency under different parameters was observed using fluorescence microscopy. Plasmids and cells were mixed in a 100 μl electroporation tip and subjected to electroporation. The transfected cells were then transferred to 6 cm culture dishes and incubated for 48 hours.

After 48 hours, the cells were trypsinized and resuspended in the flow cytometry buffer. Cells displaying green fluorescence were selected using the BD FACSMelody cell sorter (BD Biosciences, NJ, USA) and seeded at a single cell per well in a 96-well plate. The cell plate was placed in a cell culture incubator and cell growth was observed and recorded daily using a microscope. Once the wells were full, the cell colonies derived from single-cell culture were collected and transferred to larger culture plates for further expansion.

Once the cell colonies reached a sufficient size, a portion of the cells was digested using the QIAprep&amp CRISPR Kit (Qiagen) according to the product manual. PCR experiments were performed using the Mastercycler nexus gradient thermal cycler (Eppendorf, Hamburg, Germany) to amplify the target regions. PCR products with the correct molecular weight were selected and purified using the QIAquick PCR Purification Kit (Qiagen) to remove impurities. The purified samples were then sent to the Australian Genome Research Facility for Sanger sequencing to validate the complete knockout of the target exons.

### 2.4 Colony formation assay

Cells in the logarithmic growth phase were harvested and resuspended by gentle pipetting to ensure adequate dispersion. Cells were counted and adjusted to 200 cells/ml using culture medium containing 10% fetal bovine serum. The cell suspension (1 ml) was seeded into each well of a 6-well plate, followed by adding 1 ml of complete culture medium to each well. The 6-well plate was placed in a cell culture incubator and incubated for approximately 10 days. When colonies of appropriate size appeared, the culture medium was aspirated, and the cells were gently washed with PBS. The cells were then fixed and stained by incubating with a cell staining solution containing 6% paraformaldehyde and 0.5% crystal violet for 30 minutes. After careful removal of the staining solution, the cells were air-dried at room temperature. The colonies containing more than 50 cells in each well of the 6-well plate were counted and recorded.

### 2.5 Cell migration assay

In this experiment, wound closure rate was used to measure the cell migration in a two-dimensional setting. Cells were seeded in a 96-well plate and cultured until they reached 90% confluence. The culture medium then was replaced by media containing a low serum concentration and the mitotic inhibitor 5-fluoro-2’-deoxyuridine (FUDR) and treated for 12 hours. Circular or linear wounds were created at the bottom of the cell plate using a vacuum device. Then, a low dose of CKI in FUDR medium or blank FUDR medium was added. At 0 hours, the initial wound area was captured using an Olympus inverted microscope (using a 10x objective) equipped with a Canon EOS 6D camera (Canon, Tokyo, Japan). A second capture was performed between 18-24 hours to compare the wound area (ensuring that the wound had not completely closed). The images were batch standardized using XnConvert software, and the wound area was quantified using NIH ImageJ software (U.S. National Institutes of Health, MD, USA) for comparison.

### 2.6 Cell viability assays

The sensitivity of cells with different genotypes to CKI was characterized using a cell viability assay to quantify the number of live cells by producing an orange formazan dye from a water-soluble tetrazolium salt. Based on preliminary experiments, 2-3 different concentrations of CKI were selected for each cell line. Cells in the logarithmic growth phase were digested and adjusted to the desired concentration using complete medium. Then, 50 μl of the cell suspension was seeded into a 96-well plate. After overnight incubation, 50 μl of complete growth medium containing the appropriate concentration of CKI was added, and further cultured for 24 or 48 hours. To assess cell viability, 10 μl of CCK8 solution (abcam, Cambridge, UK) was added to each well and incubated for 4 hours. The absorbance was then measured at a wavelength of 450 nm using a microplate reader. Control wells without cells were included to account for background absorption. The proliferation ratio was calculated by dividing the absorbance of the wells treated with CKI by the absorbance of the control wells.

### 2.7 Cell cycle assay

We selected all knockout genotypes of the A431 cell line and CYP1A1/CYP1B1 knockout cells of the other three cell lines for validation in the cell cycle assay. Based on preliminary experiments, we treated the A431 cell line with a 1/20 dilution of CKI, E0771 cell line with a 1/15 dilution of CKI, and MDA-MB-231 and B16F1 cell lines with a 1/10 dilution of CKI. The appropriate number of cells were seeded in 1 ml of complete medium in six-well plates and incubated overnight. The following day, 1 ml of medium containing the respective CKI concentrations was added, and further incubated for 24 or 48 hours. At the endpoint of the treatment, cells were harvested, and then fixed in ice-cold ethanol solution. The fixed cells were washed with PBS after ethanol and then treated with DNA Extraction Buffer for five minutes. Subsequently, the cells were stained with propidium iodide (PI) solution containing ribonuclease for 30 minutes. The stained cells were analyzed using the LSRFortessa flow cytometer (BD Biosciences, NJ, USA) and the data were processed and analyzed using FlowJo software (TreeStar Inc., OR, USA).

### 2.8 Materials

T4 DNA Ligase kit (Cat. M0202), Adenosine 5’-Triphosphate (ATP, Cat. P0756), T4 Polynucleotide Kinase Reaction Buffer (Cat. B0201), NEBuffer 2.1 (Cat. B7202) and BbsI (Cat. R0539) were purchased from New England Biolabs, Ipswich, Ma, USA. DL-Dithiothreitol solution (Cat. 646563), Agarose (Cat. A9539), Propidium iodide solution (Cat. P4864), 6% glutaraldehyde (Cat. 340855), 0.5% crystal violet (Cat. V5265), Ribonuclease A from bovine pancreas (Cat. R5613), 5-fluoro-2′-deoxyuridine (Cat. F0503), Bovine serum albumin (Cat. A7030) and TMB solution (Cat. T0440) were purchased from Sigma-Aldrich, St. Louis, MO, USA. DH5α competent cells (Cat. 18265017) and Dulbecco’s phosphate buffered saline (Cat. 14040133) were purchased from Thermo Fisher Scientific, Waltham, MA, USA. 200bp DNA Ladder (Cat. GWS-DMW-200) were purchased from GeneWorks, Thebarton, SA, AU. Midori Green Direct (Cat. MG06) were purchased from Nippon Genetics EUROPE, Düren, Germany.

### 2.9 Statistical analysis

All experiments were repeated three times with three technical replicates (N=3). All statistical analyses were performed using GraphPad Prism version 9.5.1 for Mac OS X (GraphPad Software, San Diego, California USA). Unpaired student t test was used to compare wild cells to knockout genotypes and * was used to label the P value (* P<0.05; **P<0.01; ***P<0.001; ****P<0.0001).

## 3 Results

### 3.1 Preparing knockout genotypes with the exon dropout method

In the conventional CRISPR-Cas9 gene knockout method, an approximately 20 bp segment in the target gene’s exon is typically deleted, causing a frameshift mutation and resulting in the complete inactivation of the gene. However, it can be challenging to select appropriate deletion fragments due to the length of exons or the consideration of different splice isoforms. In this study, we used the pDG458 plasmid provided by Professor Paul Thomas at the South Australian Genome Editing Center and Genome Editing Laboratory at SAHMRI. This plasmid contains two independent exogenous DNA integration sites, allowing for the simultaneous introduction of two guide sequences^11,12^. For each target gene, we designed guide sequences on both sides of the exons to induce the complete dropout of the entire exon from the genome and to cause frameshift mutations in downstream exons, thereby completely inactivating the target gene (Figure 1a). Additionally, due to the longer deleted sequences, it was easier to perform initial screening of successfully knocked-out cell lines using PCR followed by confirmation of the knockout results through Sanger sequencing. We present our results below, in two parallel streams, one for A431 cells, where more comprehensive experiments have been carried out, and one for the other cell lines.

**Figure 1.**
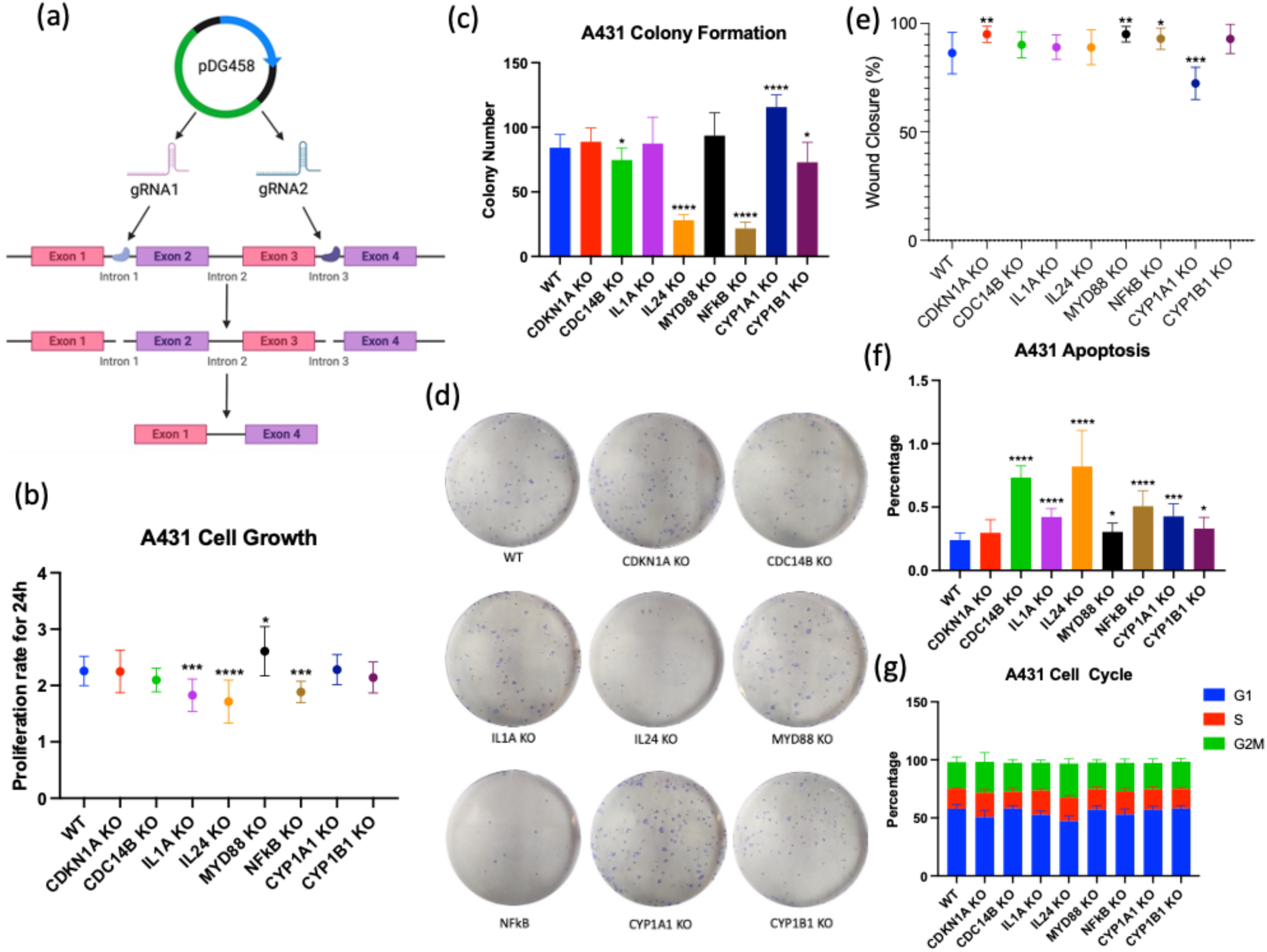
A simplified workflow for gene knockout by exon drop-out and the phenotypic changes in A431 cells after knocking out the target genes. (a) Schematic representation of the gene knockout process using the pDG458 plasmid via the exon dropout method. (b) Comparison of the 24-hour cell growth rates determined by cell viability assays between different knockout cell lines derived from the A431 cell line and the original cell line. (c) Comparison of colony formation numbers between different knockout cell lines derived from the A431 cell line and the original cell line. (d) Comparison of colony formation experiment results among different knockout cell lines derived from the A431 cell line and the original cell line (photos). (e) Comparison of cell migration rates by analysing wound closure ratios between different knockout cell lines derived from the A431 cell line and the original cell line. (f and g) Comparison of apoptotic cells and the proportions of cells in different cell cycle phases between different knockout cell lines derived from the A431 cell line and the original cell line. Results are presented as means ± SD (N = 3). Statistical analyses were conducted by comparing different knockout cell lines to the wild type (WT) (*p < 0.05, **p < 0.01, ***p < 0.001, ****p < 0.0001) using an Unpaired Student’s t-test.

### 3.2 Phenotype changes after knockout in A431 cells

To elucidate the impact of gene knockout on various cellular activities, we designed several experiments including the following assays: cell viability, colony formation, propidium iodide staining with flow cytometry, and cell migration to assess the effects of gene knockout on different aspects of cellular behaviour.

In the cell viability assay, the differences in cell proliferation ratios after 24 hours indicated that knockout of IL1A, IL24, and NFkB genes significantly reduced the proliferative activity of A431 cells. However, knockout of the MYD88 gene enhanced cell proliferation (Figure 1b). The results of the colony formation assay revealed that knockout of CDC14B, IL24, NFkB, and CYP1B1 genes significantly reduced the ability of single cells to form colonies (Figure 1c). This decrease was particularly prominent in the IL24 and NFkB knockout strains, which can be clearly observed in the images of the fixed-stained six-well plates (Figure 1d). On the other hand, knockout of the CYP1A1 gene resulted in a significant enhancement of colony formation capacity. Furthermore, cell cycle analysis revealed that, except for CDKN1A, knockout of other genes significantly increased the proportion of apoptotic cells, with IL24, NFkB, and CDC14B knockout genotypes showing more pronounced increases, consistent with the results of the cell viability assay (Figure 1f). The representation of cell proportions in different cell cycle phases also indicated that knockout genotypes like IL24 had a decreased proportion in the G1 phase and an increased proportion in the G2/M phase, suggesting cell cycle arrest (Figure 1g). In the cell migration assay, it was observed that the A431 cell line exhibited high migration efficiency, with the ability to almost cover the wound area completely within 24 hours (Figure 1e). Except for the knockout of the CYP1A1 gene, which significantly reduced cell migration speed, other knockout genotypes showed a slight increase in cell migration speed or had minimal impact compared to the wild-type cells.

### 3.3 Phenotype changes after knockout in MDA-MB-231, E0771 and B16F1 cells

The same experiments were performed on the remaining three cell lines. It can be observed that there were significant phenotypic differences in the knockout strains of the respective cell lines (Figure 2). In the MDA-MB-231 cell line, knockout of cell cycle pathway-related genes CDKN1A and CDC14B resulted in a significant decrease in the number of cell colonies formed (Figure 2a). Knockout of CYP1A1 and CYP1B1 genes led to reduced cell growth and migration rate (Figure 2c & f). In the B16F1 cell line, which has a fast growth rate and low migration rate, the knockout of genes had a relatively smaller impact on cell phenotypes. Knockout of the MYD88 gene significantly improved the colony formation capacity of B16F1 cells, while knockout of the CYP1B1 gene accelerated cell growth and migration (Figure b, d & g). Due to the incomplete adhesion of E0771 cell line, it was not possible to perform the colony formation assay as it requires washing and staining. However, an interesting finding is that the knockout of all eight genes significantly reduced the migration efficiency of E0771 cells (Figure 2h).

**Figure 2.**
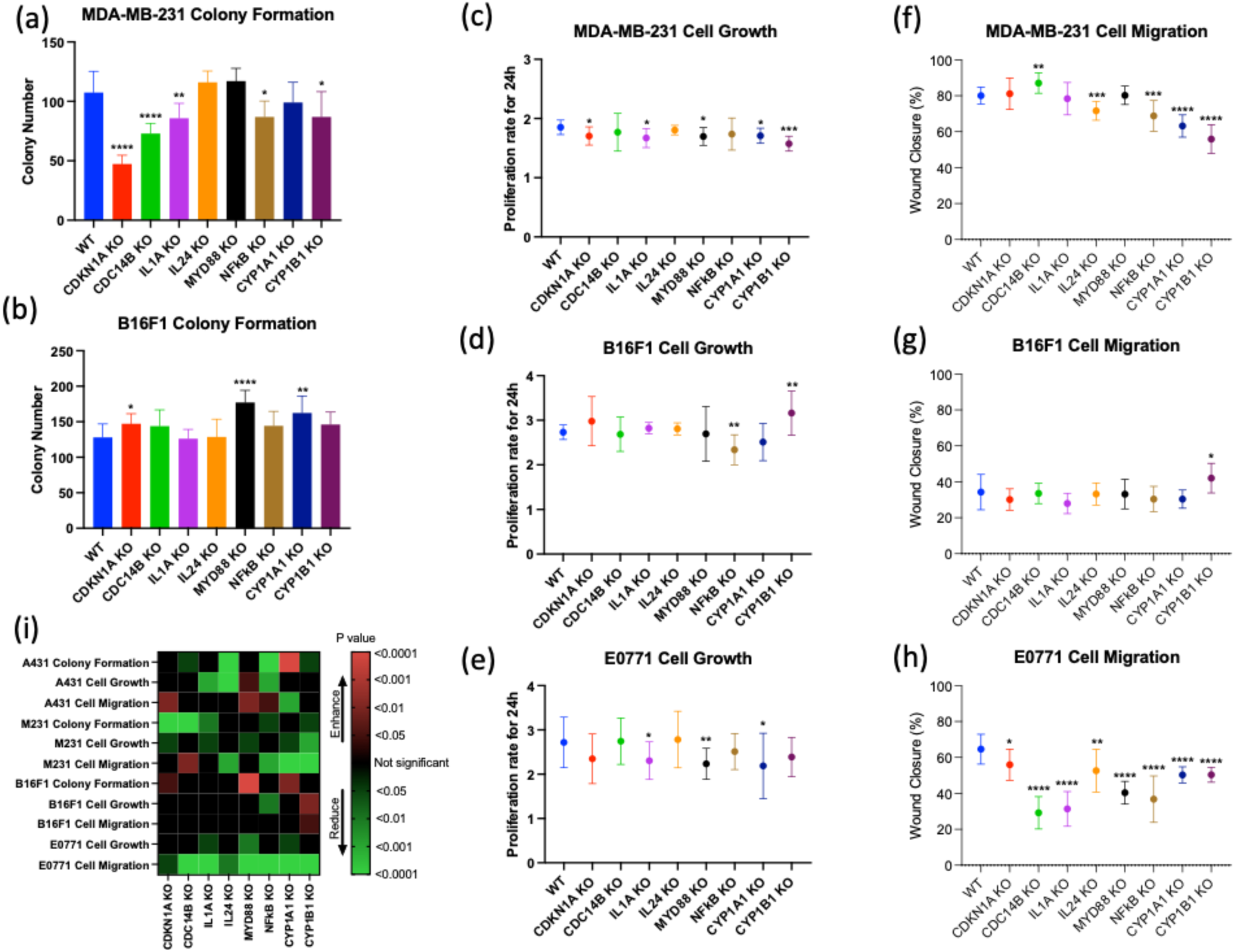
Phenotypic changes in MDA-MB-231, B16F1, and E0771 cells after gene knockout. (a and b) Comparison of colony formation numbers between different knockout cell lines of MDA-MB-231 and B16F1 cells and the original cell lines. (c, d, and e) Comparison of the 24-hour cell growth rates measured by cell viability assays between different knockout cell lines of MDA-MB-231, B16F1, and E0771 cells and the original cell lines. (f, g, and h) Comparison of cell migration rates by wound closure ratio between different knockout cell lines of MDA-MB-231, B16F1, and E0771 cells and the original cell lines. (i) A heatmap summarizing the changes in various phenotypic experiments of different cell lines compared to the original cell lines. Red indicates enhanced cell viability, green indicates reduced cell viability, and the depth of colour indicates the magnitude of the p-values for the associated changes. Results are presented as means ± SD (N = 3). Statistical analyses were performed by comparing different knockout cell lines to the wild-type (WT) using Unpaired Student’s t-test (*p<0.05, **p < 0.01, ***p <0.001, ****p <0.0001).

Based on all phenotype results, we generated a heatmap (Figure 2i) to summarize the phenotypic changes in the knockout strains of each cell line. From the heatmap, it can be observed that, overall, knockout of these genes tends to weaken cell activity in most cases. The B16F1 cell line appears to be less sensitive to gene loss, while the E0771 cell line shows a significant impact on cell migration ability due to gene knockout. Interestingly, the loss of the MYD88 gene results in enhanced cell activity in several phenotypic aspects.

### 3.4 Effect of different gene deletions on the actions of CKI in A431 cells

To test the impact of different gene deletions on the effect of CKI, we first used the CCK8 assay to evaluate cell viability under various concentrations of CKI in different cell lines. The bar graph illustrates the relative cell viability of each gene type under different CKI concentrations compared to the control group without treatment, with lower bars indicating a more pronounced effect of CKI. In the A431 cell line, almost all knockout strains showed significant differences compared to the wild type under the influence of CKI. These differences were more pronounced with longer exposure times and higher concentrations of CKI. Most of the knockout strains exhibited a weakened effect of CKI, suggesting that the function or associated products of these genes may be involved in mediating the effects of CKI. However, the deletion of MYD88 and NFkB resulted in an enhanced response to CKI, indicating that these genes may suppress the effects of CKI. Nevertheless, no single gene knock out could completely reverse the effects of CKI (Figure 3a).

**Figure 3.**
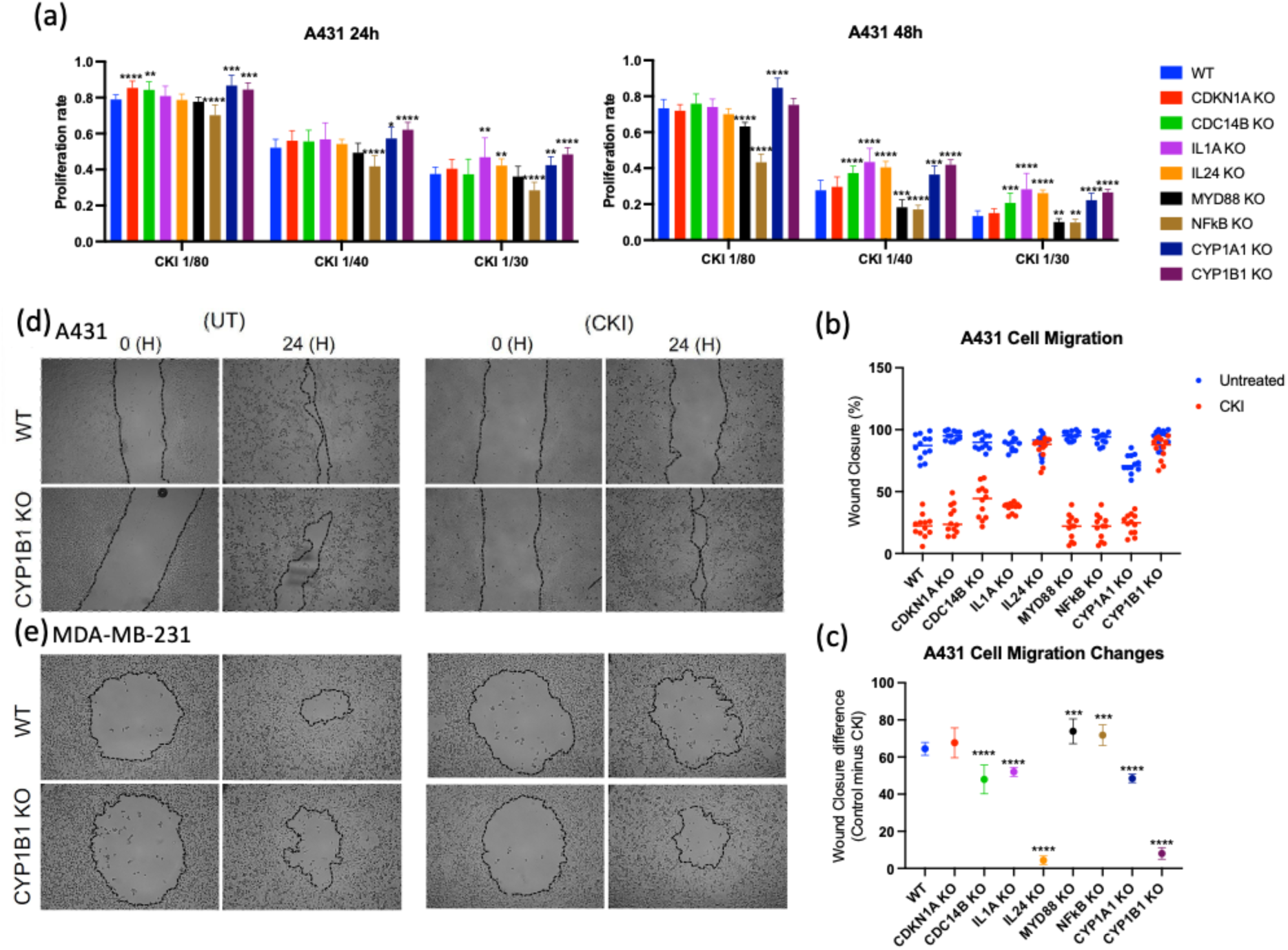
The impact of gene knockout on CKI efficacy in A431 cells. (a) The change in CKI efficacy in A431 cell lines between the original cell line and different knockout cell lines under different CKI concentrations and durations of treatment (left: 24h, right: 48h) as determined by cell viability assays. Different bars represent the ratio of cell viability resulting from CKI treatment of the respective genotype compared to the control. (b) A comparison of cell migration rates between the original A431 cell line and different knockout cell lines under CKI treatment (in red) and control (in blue) by wound closure ratio. (c) A comparison of changes in cell migration rates under CKI treatment between the original A431 cell line and different knockout cell lines. (d and e) A comparison of cell migration rates between the original cell lines and CYP1B1 knockout cell lines in A431 and MDA-MB-231 cell lines under CKI treatment and control. Results are presented as means ± SD (N = 3). Statistical analyses were performed by comparing different knockout cell lines to the wild-type (WT) using Unpaired Student’s t-test (*p<0.05, **p <0.01, ***p <0.001, ****p <0.0001).

In a previous study, we found that CKI significantly reduces the migration rate of tumor cells. Therefore, in this study, we conducted experiments to compare the effects of CKI on cell migration in different gene knockout cells. In the A431 cell line, the comparison of wound healing rates between CKI treatment and the control group in different gene knockout cells clearly showed that CKI significantly decreases cell migration speed (Figure 3b). However, in the IL24 and CYP1B1 gene knockout cells, this difference is mostly eliminated, indicating the importance of these two genes in mediating the effect of CKI on cell migration in A431 cells. This difference can also be clearly observed in the comparison of cell images of wound healing (Figure 3d). Further comparison of the differences in cell migration speed between control group and CKI treatment group in different genotypes showed that the knockout of the MYD88 and NFkB genes enhances the inhibitory effect of CKI on cell migration, while the knockout of the other genes, except CDKN1A, significantly weakens the effect of CKI (Figure 3c).

### 3.5 Effect of different gene deletions on the actions of CKI in MDA-MB-231, E0771 and B16F1 cells

Similar screening experiments were conducted on the other three cell lines (Figure 4). It can be observed that these cell lines exhibited less pronounced differences in their response to CKI after gene knockout compared to A431. Most of them showed a decrease in CKI activity, except for the knockout of the CYP1B1 gene in the B16F1 cell line and the knockout of the IL1A gene in the E0771 cell line, which exhibited an enhanced response to CKI (Figure 4a, b & c). The heatmap (Figure 5a) provides a visual summary of the results from the cell viability assays. It can be observed that most of the gene knockout strains exhibited a reduction in the effects of CKI. The changes were more pronounced in the A431 cell line, while the MDA-MB-231 and B16F1 cell lines showed smaller changes in response to CKI after gene knockout. Additionally, the deletion of the CYP1A1 and CYP1B1 genes resulted in altered cellular responses to CKI in each cell line.

**Figure 4.**
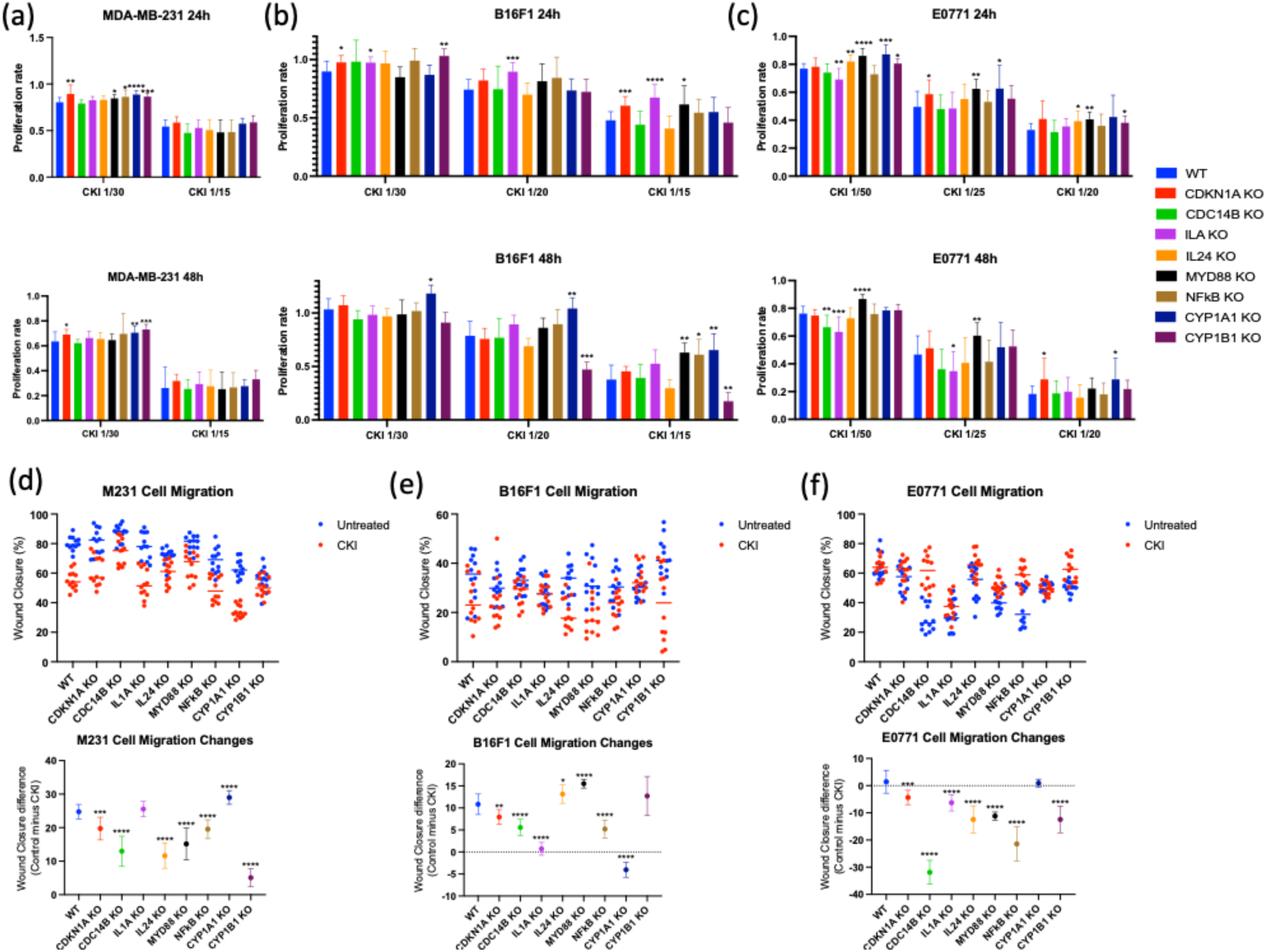
The impact of gene knockout on CKI efficacy in MDA-MB-231, B16F1, and E0771 cells. (a, b, and c) CKI efficacy change in MDA-MB-231, B16F1, and E0771 cell lines between the original cell line and different knockout cell lines under different CKI concentrations and treatment durations (top: 24h, bottom: 48h) as determined by cell viability assays. Different bars represent the ratio of cell viability under CKI treatment of the respective genotype compared to the control. (d, e, and f) A comparison of cell migration rates between the original MDA-MB-231, B16F1, and E0771 cell lines and different knockout cell lines under CKI treatment (in red) and control (in blue) by wound closure ratio (top); a comparison of changes in cell migration rates under CKI treatment between the original cell lines and different knockout cell lines (bottom). Results are presented as means ± SD (N = 3). Statistical analyses were performed by comparing different knockout cell lines to the wild-type (WT) using Unpaired Student’s t-test (*p<0.05, **p <0.01, ***p <0.001, ****p <0.0001).

**Figure 5.**
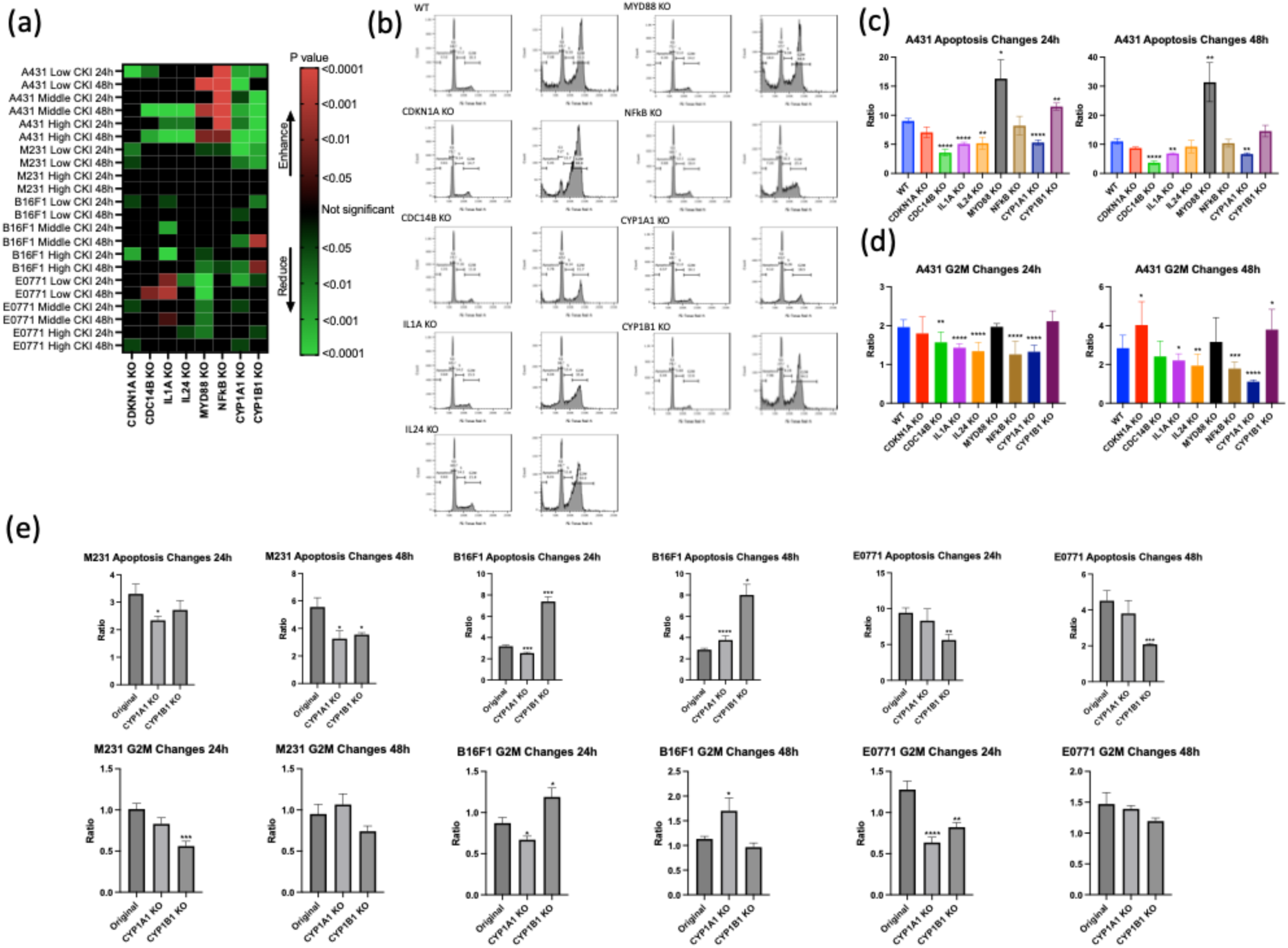
Summary of cell viability experiments and determination of CKI efficacy in cell cycle experiments. (a) A heatmap summarizing the change in CKI efficacy for different genotypes in cell lines as observed in cell viability experiments. Red indicates enhanced CKI efficacy, while green represents reduced CKI efficacy, with the depth of colour reflecting the p-value’s magnitude of the relevant changes. (b) Representative histograms illustrating cell cycle phases for different genotypes with or without CKI treatment in A431 cells. (c) A comparison of the fold increase in apoptosis rates under CKI treatment for different A431 knockout cell lines. (d) A comparison of the fold increase in the percentage of G2M phase cells under CKI treatment for different A431 knockout cell lines. (e) A comparison of the fold change in the percentage of apoptotic cells and G2M phase cells under CKI treatment for CYP1A1 and CYP1B1 knockout cell lines in MDA-MB-231, B16F1, and E0771 cells compared to the original cell lines. Results are represented as means ± SD (N = 3). Statistical analyses were performed by comparing different knockout cell lines to the wild-type (WT) using Unpaired Student’s t-test (*p<0.05, **p <0.01, ***p <0.001, ****p <0.0001).

The effects of CKI on reducing cell migration speed in the MDA-MB-231 and B16F1 cell lines were not as significant as in the A431 cell line (Figure 4d & e), but it was still clear that the deletion of most genes weakens the effect of CKI. In the MDA-MB-231 cell line, like A431, the knockout of IL24 and CYP1B1 genes had a significant impact on the effect of CKI, while the knockout of CYP1A1 gene enhanced the effect of CKI (Figure 3e & 4d). In the B16F1 cell line, except for the enhancement of CKI’s inhibitory effect on cell migration by the knockout of IL24 and MYD88, the deletion of other genes generally weakened the effect of CKI (Figure 4e). The results in the E0771 cell line are quite unique, as multiple gene knockout cells show an increase in cell migration efficiency upon CKI treatment (Figure 4f). Due to the loose adherence of this cell line, we observed a significant increase in floating cells and cell clusters in the CKI-treated samples during the experiment. Therefore, the apparent migration increases may be due to the floating cells reattaching to the wound area, resulting in a phenomenon that could be misinterpreted as migration but reflects only random dispersal. This effect should be investigated in future experiments using 3D cell migration assays to eliminate this influence.

### 3.6 The changes in cell cycle with CKI treatment

We then selected all gene knockout strains of the A431 cell line and knockout strains of the CYP1A1 and CYP1B1 genes in other cell lines for subsequent propidium iodide staining experiments. Figure 5b shows the cell cycle profiles of all different gene knockout strains of the A431 cell line under CKI treatment compared to the control group. Although there were minimal differences in the morphology of the cell cycle profiles among the knockout strains under control conditions, noticeable differences were observed under the influence of CKI. Apart from a significant difference in the proportion of apoptotic cells, there were significant variations in the overall cell cycle profiles, particularly in the proportion of cells in the G2/M phase. For example, a comparison between the CDKN1A knockout strain and the CYP1A1 knockout strain indicated that the cells underwent cell cycle arrest in the G2/M phase. The changes in frequencies of apoptotic cells and G2/M phase among different gene knockout strains under CKI treatment were compiled, and showed that in the A431 cell line, the knockout of the MYD88 gene and the CYP1B1 gene significantly enhanced the effect of CKI, while the knockout of other genes generally weakened the effect of CKI (Figure 5c & d). The knockout strains of the CYP1A1 and CYP1B1 genes in the other cell lines were also analyzed for changes in the proportion of apoptotic cells and G2/M phase under CKI treatment (Figure 5e), and found to show consistency with cell viability results.

Additionally, a comparison to the results of the cell viability assays is shown in Tables 1 and 2. It can be observed that the results of the cell cycle analysis aligned with the results of the cell viability assay. Knockout of the CYP1A1 gene reduced the effectiveness of CKI, while the loss of CYP1B1 function enhanced the effect of CKI in the B16F1 cell line but reduced it in the other two cell lines. We also summarized the results of CKI in the A431 cell line, where significant differences were observed among different genotypes (Table 2). The knockout of the CDKN1A gene had the smallest impact, while the deletion of MYD88 and NFkB enhanced the effect of CKI in various aspects. The knockout of the remaining genes resulted in a decrease in the effect of CKI.

**Table 1.**
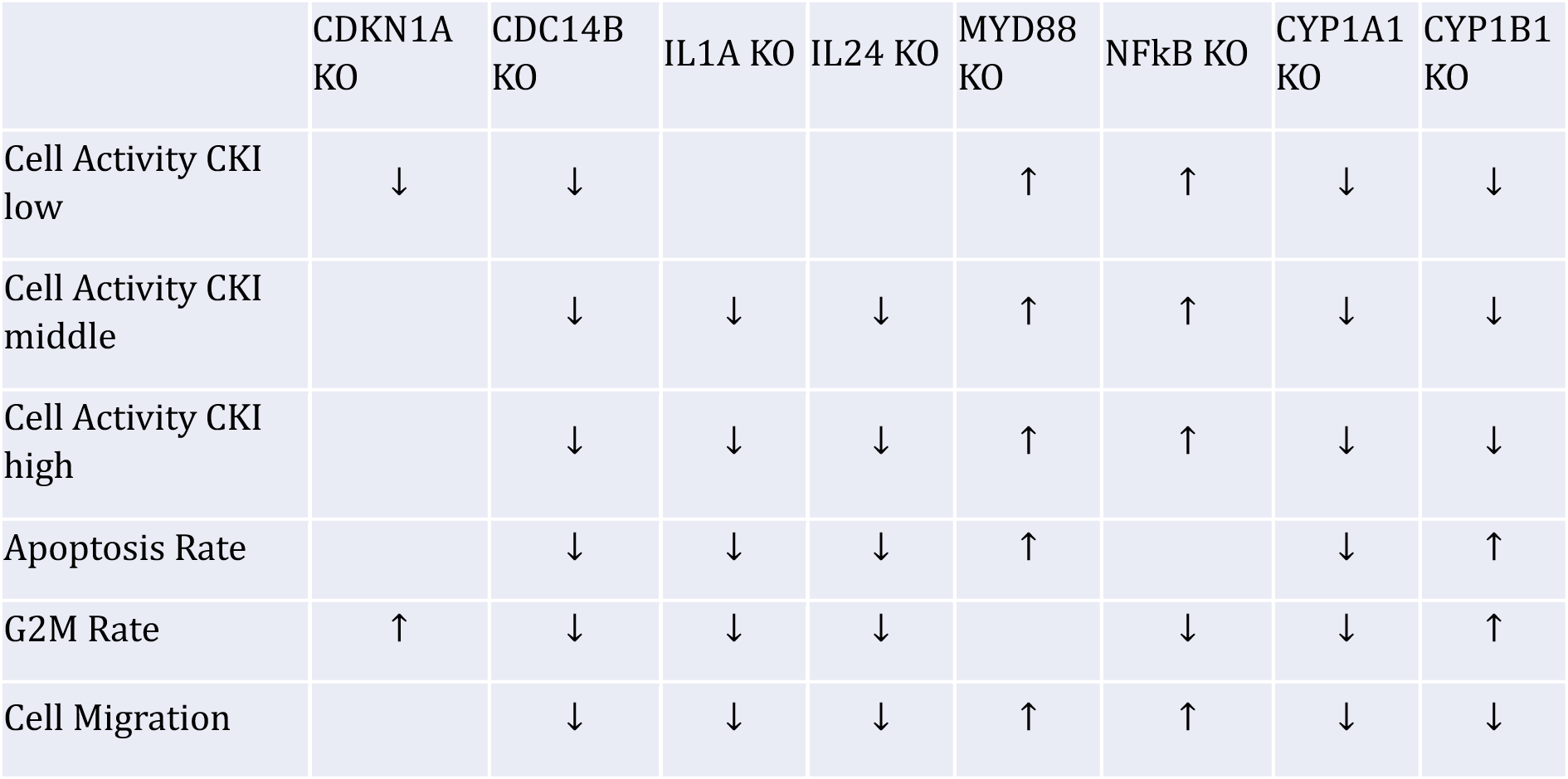
Summary of the changes in sensitivity to CKI of A431 cell lines with different gene knockouts in various experiments (compared to the original cell line).↑ indicates enhanced CKI efficacy, and ↓ indicates reduced CKI efficacy.

**Table 2.**
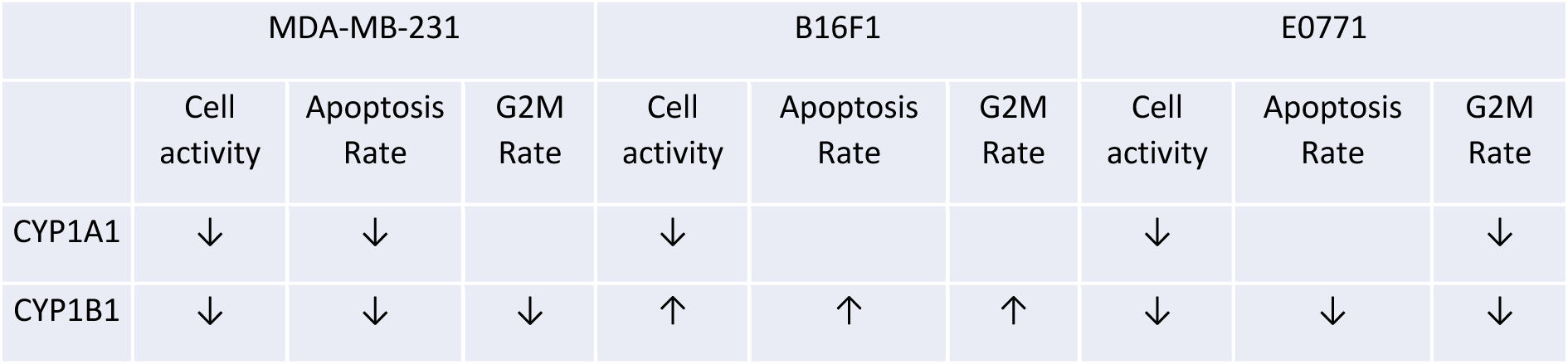
Summary of the changes in sensitivity to CKI of MDA-MB-231, B16F1, and E0771 cell lines with CYP1A1 and CYP1B1 knockouts in various experiments (compared to the original cell line).↑ indicates enhanced CKI efficacy, and ↓ indicates reduced CKI efficacy.

In summary, we successfully established knockout cell lines for 8 genes in four different cell lines using the exon dropout method and elucidated their phenotypic changes through various experiments. Furthermore, by observing how the effects of CKI changed after the knockout of different genes, we confirmed that these genes and their associated signalling pathways significantly influence CKI’s anti-tumor activity. However, it’s important to note that the knockout of any single gene was unable to completely counteract its pharmacological effects. This result further underscores the proposed mechanism of multi-target synergistic action in traditional complex mixtures of natural products such as Chinese medicine formulas.

## 4 Discussion

Different types of tumor cells often exhibit significant differences in their molecular genetics and receptor expression profiles, which in turn influence their physiological activities and sensitivity to therapeutic interventions^14, 15^. In this project, there were substantial variations in sensitivity to CKI, phenotypic changes after different gene knockouts, and CKI’s pharmacological effects among the various cell lines. In the case of the two human cell lines, MDA-MB-231 cells which are classified as triple-negative breast cancer, demonstrated lower sensitivity to the drug compared to A431 cells. The experiments required CKI concentrations more than two-fold higher than those used for the A431 cell line, which overexpresses the epidermal growth factor receptor, to achieve similar antitumor effects^16^. It’s also evident that in the A431 cell line, the effects of different knockout cell strains on CKI’s antitumor action were significant and consistent. This emphasizes the critical roles of heterogeneous genes and pathways in CKI’s antitumor pharmacology. In the case of MDA-MB-231, the impact of gene knockout on CKI’s efficacy was not as pronounced, but the results that did show significant impact were consistent with those observed in A431 cells. However, in the two mouse-derived cell lines, apart from their reduced sensitivity compared to A431 cells, there were discordances with the results obtained from human-derived cell lines. These variations indicate the need for special consideration of the differences between different cell lines, especially those from different sources, in complex compound pharmacological experiments, such as traditional Chinese medicine.

Due to the poorly restrained proliferation characteristic of tumor cells, genes related to the cell cycle pathway have always been important subjects in cancer research and the study of anti-cancer drug targets. The CDKN1 gene encodes the P21 protein, which acts as a regulator of cell cycle progression at G1^17,18^. It is often deregulated in human cancer. Depending on the cellular context, P21 can either promote or inhibit tumorigenesis^19^. The knockout of this gene significantly inhibited cell growth and colony formation in the MDA-MB-231 cell line. Another gene that was knocked out is CDC14B, which is involved in the exit of cell mitosis and initiation of DNA replication. CDC14B overexpression can induce oncogenic transformation, and it has been shown to interact with and dephosphorylate the tumour suppressor protein P53^20^. Knocking out this gene reduced colony formation in both A431 and MDA-MB-231 cells and increased the proportion of apoptotic cells. Additionally, the cell cycle pathway is one of the major pathways through which CKI exerts its cytotoxic effect, leading to significant cell cycle arrest when acting on tumor cells. Transcriptome analysis indicated that CKI generally downregulates genes in the cell cycle pathway^3^. However, in this experiment, the selected genes, CDKN1A and CDC14B, showed significantly upregulated expression under the influence of CKI. Although the loss of CDKN1A had a less noticeable impact on CKI’s effects, knocking out the CDC14B gene consistently weakened the effects of CKI in all experiments with A431 cell lines and other cell lines in cell migration assays. This suggests that CDC14B may be a critical node gene for CKI to exert its pharmacological effects within the cell cycle pathway.

Cytokines play a significant role in the occurrence, development, and treatment of tumours. Additionally, cytokines and their related receptor pathways are among the pathways in which CKI’s regulatory effect is very pronounced, as shown in our previous studies^6, 21^. IL-1 signalling induces the expression of inflammatory mediators, including IL-6 and COX2, which promote the survival and proliferation of malignant cells^22^. After knocking out IL1a, we observed a slowing of tumour cell growth and a reduction in CKI’s effects in skin cancer cells. However, this effect may not be very pronounced, possibly due to compensatory actions by exogenous IL1b. IL24 is a broad-spectrum anticancer gene capable of inducing apoptosis or selective toxic autophagy in tumour cells^23^. After knocking out the IL24 gene, a significant reduction in cell growth and colony formation in the A431 cell line was observed, along with a weakening of CKI’s effects. Interestingly, CKI’s inhibition of cell migration was entirely offset in this knockout strain. Given the reports that IL24 can inhibit cancer cell invasion and migration, this suggests that CKI’s stimulation of IL24 production might be a key factor in inhibiting cell migration^24^.

The NF-kB gene encodes a subunit of the transcription factor complex known as nuclear factor-kappa-B, which acts as a central activator of genes involved in inflammation and immune function. MYD88 serves as a critical signal transducer in the interleukin-1 and Toll-like receptor signalling pathways, and its effects are largely reliant on NF-kB^25,26^. When NF-kB signalling is activated, it can render cells resistant to chemotherapeutic drugs and inhibit cell apoptosis. Therefore, the upregulation of these two genes in response to CKI treatment may represent a self-protective mechanism in cells against the cytotoxic effects of CKI. Consequently, after the knockout of these two genes, the effects of CKI were significantly enhanced, as demonstrated in the summary of changes in CKI responses following gene knockouts in the A431 cell line. MYD88, as a crucial nodal gene linking cytokines and intracellular responses, had previously been found to play a vital role in CKI’s effects^27^. This suggests that various cytokine-mediated immune-related pathways are also important targets of CKI’s action. The results, which show enhanced anticancer effects following the knockout of MYD88, indicate that CKI mainly mediates cytokine changes in tumour cells for cell protection. Nevertheless, further studies are required to comprehensively evaluate CKI’s overall effects on immune cells and the entire organism.

As a complex mixture containing hundreds of components, CKI induces the upregulation of various metabolism-related genes in response to drug treatment. In this study, we chose to knock out CYP1A1 and CYP1B1 genes, which were identified as significantly upregulated by CKI in previous transcriptome results. Earlier research has shown that CYP1A1 typically is significantly downregulated, while CYP1B1 is markedly upregulated in human tumour cells^28,29^. These changes may be associated with the influence of the proinflammatory microenvironment on cells, which affects tumour cell physiological functions to some extent. However, interestingly these effects were not uniform across different cell lines. The significant upregulation of these genes in the presence of CKI may be due to the activation of the AhR receptor by relevant small-molecule ligands, although further validation is needed^30,31^.

After the knockout of both genes, the effects of CKI were significantly reduced, and this phenomenon was consistent across multiple experiments in human A431 and MDA-MB-231 cell lines. This suggests that the metabolic products of CYP1A1 and CYP1B1 may possess substantial pharmacological activity. Interestingly, the results indicate that while the knockout of both metabolic enzyme genes has some weakening effect on most of CKI’s pharmacological actions, the knockout of CYP1A1 essentially abolishes CKI’s G2M phase arrest effect. Conversely, the knockout of CYP1B1 does not affect this function. Furthermore, the knockout of CYP1B1 almost completely counteracts CKI’s reduction of tumour cell migration, whereas the knockout of CYP1A1 does not have a pronounced effect on this function. This suggests that the metabolic products of CYP1A1 and CYP1B1 may be key active components of CKI, responsible for inducing cell cycle arrest and preventing tumour cell migration, respectively.

In conclusion, this study building on previous transcriptomic analysis results, focused on eight selected key genes from four pathways, and establish a panel of genetic knockouts in four different cell lines. The experimental results showed that the knockout of most of the selected genes has a significant impact on the pharmacological effectiveness of CKI, thereby validating the earlier gene analysis results and the proposed mechanism of action of multi-target synergy of complex components of traditional Chinese medicine. Additionally, we discovered that IL24 and CYP1B1 genes appear to play a crucial role in CKI’s inhibition of cell migration, while CYP1A1 is important for CKI’s effect in causing G2M cell cycle arrest. The demonstration here that knockout of these genes almost completely negates the effects of CKI provides a promising starting point for further research on therapeutic strategies tailored to specific cancer types.

## Author Contributions

Conceptualization, HS and DA; investigation, HS, SN, YZ; formal analysis, HS, ZQ and YH-L; resources, WW; data curation, HS; writing-original draft preparation, HS; writing-review and editing, DA and AY; supervision, DA and AY.

## Acknowledgments

The authors would like to thank Michaela Scherer and Jayshen Christa Arudkumar for technical support in CRISPR/Cas process.

## Abbreviations

CRISPR/CAS: Clustered regularly interspaced short palindromic repeats and CRISPR associated protein
CDKN1A: Cyclin dependent kinase inhibitor 1A (ENSG00000124762)
CDC14B: Cell division cycle 14B (ENSG00000081377)
IL1A: Interleukin 1 alpha (ENSG00000115008)
IL24: Interleukin 24 (ENSG00000162892)
MYD88: Myeloid differentiation primary response 88 (ENSG00000172936)
NFkB: Nuclear factor kappa-light-chain-enhancer of activated B cells (ENSG00000109320)
CYP1A1: Cytochrome P450 family 1 subfamily A member 1 (ENSG00000140465)
CYP1B1: Cytochrome P450 family 1 subfamily B member 1 (ENSG00000138061)

