## Supplementary Figure 1 and Table 1 for "Validating the Target Functions and Synergistic Multi-Target, Multi-Pathway Action Mode of Compound Kushen Injection Using CRISPR/CAS"

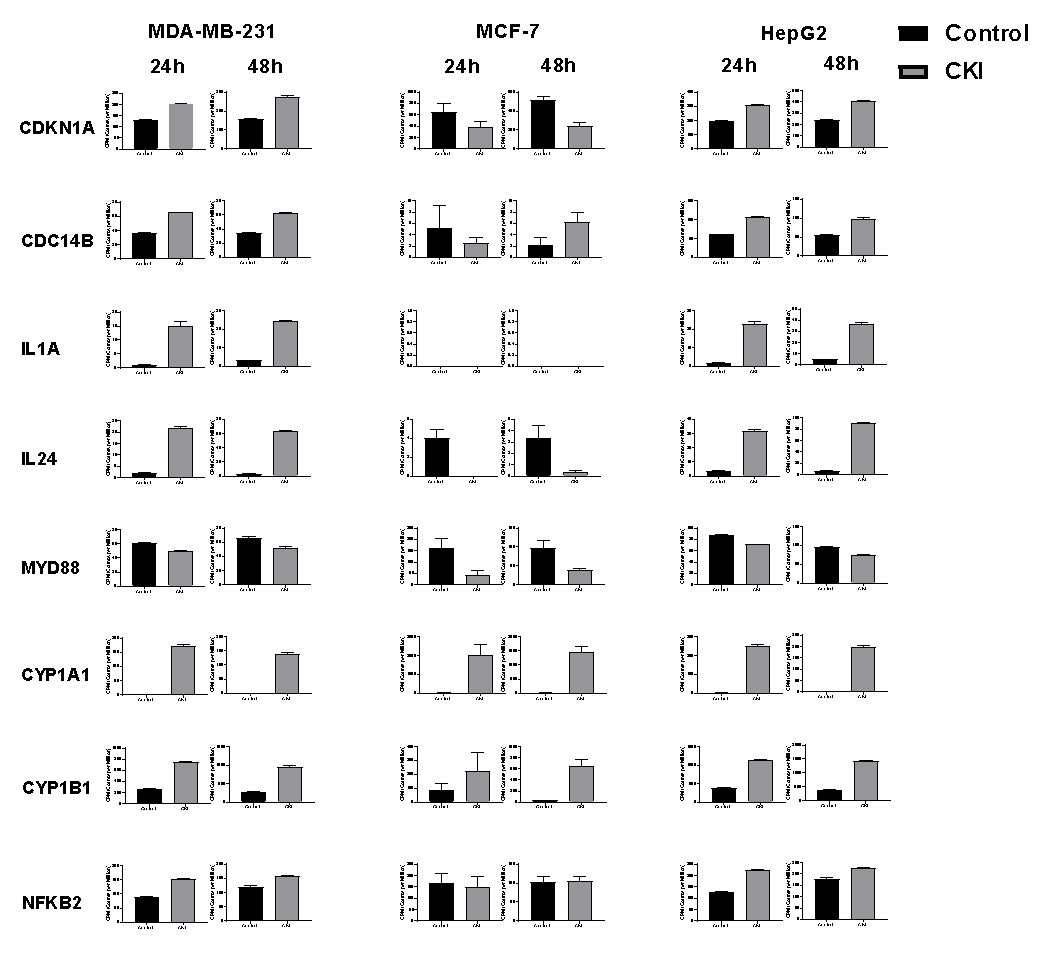

Supplementary Figure 1. Expression level (counts per million) of selected genes in our previous sequencing data^3,4,21^. Results are represented as means ±SEM (n=3)

|  | HUMAN | MOUSE |
| --- | --- | --- |
| CDC14B | Guides  1.  CACCGTCATGCACGTTTCAGATGGT  AAACACCATCTGAAACGTGCATGAC  2.  ACCGGTTATAAATGGGATGGCCTGT  TAAAACAGGCCATCCCATTTATAAC  PCR Primers  FWD:5'-GCTAGCCATCACATGCATGCTGC-3' | Guides  1.  CACCGCTGCCATCTGAAACATGGGG  AAACCCCCATGTTTCAGATGGCAGC  2.  ACCGAGGTTAGTTATTCTACGATGT  TAAAACATCGTAGAATAACTAACCT  PCR Primers  FWD:5'-GCTAGCCATCACATGCATGCTGC-3' |
|  | REV:5'-TTGAGTGGTCCAAAATCTGCGT-3' | REV:5'-TTGAGTGGTCCAAAATCTGCGT-3' |
| CDKN1A | Guides  1.  CACCGTCTAATCTCCGCCGTGACC  AAACGGTCACGGCGGAGATTAGAC  2.  ACCGTTTGAGGCCCTCGCGCTTCCGT  TAAAACGGAAGCGCGAGGGCCTCAAA  PCR Primers  FWD:5'-GAGCTAGGACCACAACCCGGGT-3' | Guides  1.  CACCGAAGAAGGACCCGGGTCACTA  AAACTAGTGACCCGGGTCCTTCTTC  2.  ACCGCCTAAATTAGTAGGACGGTAGT  TAAAACTACCGTCCTACTAATTTAGG  PCR Primers  FWD:5'-GAGCTAGGACCACAACCCGGGT-3' |
|  | REV:5'-TGACAGCGATGGGAAGGAGCCA-3' | REV:5'-TGACAGCGATGGGAAGGAGCCA-3' |
| CYP1A1 | Guides  1.  CACCGAGTTGTTGAACTACCACTGT  AAACACAGTGGTAGTTCAACAACTC  2.  ACCGGAAGAAGACTATTCCACAACGT  TAAAACGTTGTGGAATAGTCTTCTTC  PCR Primers  FWD：5'-TCAGCGCTGCTTCCCCCAAATG-3' | Guides  1.  CACCGAGGCATCAGACTGAGGGCGA  AAACTCGCCCTCAGTCTGATGCCTC  2.  ACCGGAATGTACTCTATCCTCAGCGT  TAAAACGCTGAGGATAGAGTACATTC  PCR Primers  FWD：5'-TCAGCGCTGCTTCCCCCAAATG-3' |
|  | REV: 5'-TGCCCCAGGGTCCTGCATGTAA-3' | REV: 5'-TGCCCCAGGGTCCTGCATGTAA-3' |
| CYP1B1 | Guides  1.  CACCGTTTAGCGGCCAAGGGTCGTT  AAACAACGACCCTTGGCCGCTAAAC  2.  ACCGATCGGAAACGCGGCGGCGGTGT  TAAAACACCGCCGCCGCGTTTCCGAT  PCR Primers  FWD:5'-AGCTCCGACCTCTCCACCCAAC-3' | Guides  1.  CACCGCCACTGGCCTAAGTGCACGG  AAACCCGTGCACTTAGGCCAGTGGC  2.  ACCGACGCGCCACCCGCGCAGTCGGT  TAAAACCGACTGCGCGGGTGGCGCGT  PCR Primers  FWD:5'-AGCTCCGACCTCTCCACCCAAC-3' |
|  | REV: 5'-TCCAGTGCTCCGAGTAGTGGCC-3' | REV: 5'-TCCAGTGCTCCGAGTAGTGGCC-3' |
| IL1A | Guides  1.  CACCGGGCTCCCCTCACAGATATC  AAACGATATCTGTGAGGGGAGCCC  2.  ACCGTAGGCTGTCAAAATTTTGCCGT  TAAAACGGCAAAATTTTGACAGCCTA  PCR Primers  FWD:5'-GCTAGCCATCACATGCATGCTGC-3' | Guides  1.  CACCGTTCTAATTTAGTACAGGGGC  AAACGCCCCTGTACTAAATTAGAAC  2.  ACCGTTGATTCGATAAGGGGCTCAGT  TAAAACTGAGCCCCTTATCGAATCAA  PCR Primers  FWD: 5'-ACTGTCCTCTCTCAAGGCACCT-3' |
|  | REV:5'-TTGAGTGGTCCAAAATCTGCGT-3' | REV: 5'-CCCAAGGGCTGTTTCCTCCTGG-3' |
| IL24 | Guides  1.  CACCGACGAGCGGCCTTAAGTTTGA  AAACTCAAACTTAAGGCCGCTCGTC  2.  ACCGACTCTATTCGGGACGAAGCCGT  TAAAACGGCTTCGTCCCGAATAGAGT  PCR Primers  FWD:5'-GAGCTAGGACCACAACCCGGGT-3' | Guides  1.  CACCGTATCAGGCGTGACGTTTGTT  AAACAACAAACGTCACGCCTGATAC  2.  ACCGCAGGAGGATCAATACGTCCCGT  TAAAACGGGACGTATTGATCCTCCTG  PCR Primers  FWD:5'-TGCAGGTGTGTGGAGGGAGAGG-3' |
|  | REV:5'-TGACAGCGATGGGAAGGAGCCA-3' | REV:5'-AGCAAGGTAGCTGCACCTGGGA-3' |
| MYD88 | Guides  1.  CACCGTGAGGACGACCCTCCTTCG  AAACCGAAGGAGGGTCGTCCTCAC  2.  ACCGACCGCGACCGGTTATTAAGAGT  TAAAACTCTTAATAACCGGTCGCGGT  PCR Primers  FWD：5'-TCAGCGCTGCTTCCCCCAAATG-3' | Guides  1.  CACCGTTGCTAGAATCTAGACTACG  AAACCGTAGTCTAGATTCTAGCAAC  2.  ACCGAGGATCAACGAGGCTAGCCTGT  TAAAACAGGCTAGCCTCGTTGATCCT  PCR Primers  FWD:5'-ACACCTTCGAGGGGAGGTGAGC-3' |
|  | REV: 5'-TGCCCCAGGGTCCTGCATGTAA-3' | REV:5'-ACTGGGCATCTAGCAGGGCACA-3' |
| NFKB2 | Guides  1.  CACCGGAGAGCGAGATCCGGAGTT  AAACAACTCCGGATCTCGCTCTCC  2.  ACCGCCTCCCCTAGTCCTAAAGCGGT  TAAAACCGCTTTAGGACTAGGGGAGG  PCR Primers  FWD:5'-AGCTCCGACCTCTCCACCCAAC-3' | Guides  1.  CACCGACCTTCTCTGTCTATAACGC  AAACGCGTTATAGACAGAGAAGGTC  2.  ACCGCTAGGTTCTGAGATACTGCGGT  TAAAACCGCAGTATCTCAGAACCTAG  PCR Primers  FWD:5'-ATTGGACGCTCCTCCACTCCCC-3' |
|  | REV: 5'-TCCAGTGCTCCGAGTAGTGGCC-3' | REV: 5'-TTCCGGCCCTTCTCACTGGAGG-3' |

Supplementary Table 1. The information for CRISPR guides and PCR primers for each gene.
